# Investigating lncRNAs at different levels of parasitism in parasitic plants

**DOI:** 10.1101/2024.11.26.625533

**Authors:** Wenderson Felipe Costa Rodrigues, Laura Oliveira Pires, Luiz-Eduardo Del-Bem, Juliane Karine Ishida

**Affiliations:** Graduate Program in Plant Biology, Department of Botany, Institute of Biological Sciences (ICB), Federal University of Minas Gerais (UFMG), 31270-901 Belo Horizonte, Brazil; Department of Botany, Institute of Biological Sciences (ICB), Federal University of Minas Gerais (UFMG), 31270-901 Belo Horizonte, Brazil

**Keywords:** Parasitic plants, HGT, RNA

## Abstract

Parasitic plants exhibit unique adaptations, including specialized structures like haustoria and reduced organs. We analyzed genomic and transcriptomic data from nine parasitic species across hemi-, holo-, and endoparasitic lifestyles, uncovering distinct patterns of lncRNA abundance and expression. Holoparasites exhibited the highest lncRNA counts, while endoparasites displayed notable reductions. Expression analyses revealed tissue-specific patterns, with elevated lncRNA activity in haustorial tissues of *Cuscuta*. Phylogenetic analyses suggested horizontal gene transfer (HGT) of lncRNA sequences, particularly in *Sapria*. These findings highlight the potential involvement of lncRNAs in parasitic plant adaptation, host interaction, and genomic evolution through HGT.

## Main

Long non-coding RNAs (lncRNAs) are increasingly recognized as key regulatory molecules in various organisms^1,2^. These non-coding transcripts, generally exceeding 200 nucleotides, exhibit diverse secondary structures, localization patterns, and regulatory capacities within the cell^3^. In plants, lncRNAs have been linked to the regulation of diverse developmental processes, including organ development and maturation. They are involved in critical stages such as flower development, root growth, and stress responses, often by modulating gene expression at transcriptional or post-transcriptional levels^4^. Notably, lncRNAs exhibit the ability to function at locations distant from their synthesis site, similar to other types of mobile RNA molecules, such as small interfering RNAs (siRNAs) and messenger RNAs (mRNAs)^5^. This capacity for long-distance movement suggests that lncRNAs may act as signaling molecules, coordinating developmental and physiological processes throughout the plant.

The critical roles of lncRNAs in non-parasitic plants raise questions about their potential involvement in parasitic plants, particularly given their capacity for mobility, which may facilitate complex interactions between host and parasite tissues. Parasitic plants are angiosperms with specialized adaptations that enable them to extract nutrients from host plants through structures known as haustoria^6,7^. Parasitic plants vary in their degree of dependence on the host, ranging from hemiparasites that can photosynthesize but supplement their nutrition from hosts, to holoparasites and endoparasites that rely exclusively on host-derived resources^6^. Prior studies have demonstrated that lncRNAs play essential roles in plant responses to biotic stress, including interactions with fungi or nematodes, where they contribute to pathogen resistance, gene regulation, and sexual differentiation^8,9^. These findings suggest that lncRNAs could similarly influence key aspects of parasitism, including haustorial development and function. Furthermore, beyond their role in parasitic interaction, lncRNAs are likely involved in the broader developmental processes of parasitic plants, potentially shaping the formation and differentiation of organs. Parasitic plants often exhibit modified or reduced structures, such as diminished roots or absent leaves, which are hallmark adaptations to their parasitic lifestyle^10,11^. However, despite their potential significance, no direct evidence currently links lncRNAs to these specific developmental modifications, leaving an open field for future research into their roles in both parasitism and general plant development.

*Cuscuta*, a holoparasitic genus, and *Sapria*, an endoparasitic species, exhibit notable alterations in floral growth regulation, reflecting evolutionary adaptations to their parasitic lifestyles. For instance, *Sapria* demonstrates a striking temporal mismatch in flowering relative to its host, which may represent a divergence in developmental timing mechanisms^12^. This mismatch is accompanied by low fertilization rates and the production of exceptionally large, elaborate flowers, likely as compensatory adaptations for its unique reproductive strategy. In contrast, *Cuscuta* exhibits floral growth tightly coordinated with the availability of host-derived resources, often synchronizing its reproductive phase with host availability and environmental cues, potentially mediated by host-derived hormonal signals^13^. These traits suggest a fine-tuned regulatory network in which lncRNAs might play a central role, coordinating the expression of genes involved in floral development and compensating for the extensive gene loss documented in the genomes of these parasitic plants^14,15^. Additionally, in *Cuscuta*, the bidirectional movement of lncRNAs between the parasite and its host has been identified, suggesting that these molecules may act as mediators of interspecies communication, facilitating the transfer of regulatory information crucial for parasitism^16^. However, beyond this single report, there is a significant gap in understanding the roles of lncRNAs in plant-plant parasitic interactions.

This gap highlights the need for further research to understand the regulatory potential of lncRNAs within the context of parasitism and the specific ways these molecules may be contributing to host-parasite dynamics in plants. To begin addressing this knowledge gap, we aimed to identify and characterize the lncRNAs, by leveraging genomic data and bioinformatic tools, in nine parasitic plant species with varying parasitic lifestyles, including hemi-, holo-, and endoparasites (Table S1). Our analysis revealed that there was no consistent trend in lncRNA abundance between parasitic and non-parasitic species. For instance, the basal eudicot *Aquilegia coerulea* contained approximately 4,728 lncRNAs, a number significantly higher than most parasitic species, with the exception of the holoparasitic genus *Orobanche* (Figure S1A). Species within Solanales and Lamiales, a clade that contains the parasitic plant families Orobanchaceae and Convolvulaceae, respectively, exhibited the highest lncRNA counts (Figure S1A). In contrast, the Malpighiales clade, which includes the endoparasitic Sapria himalayana, displayed a significantly lower average of around 2,035 lncRNAs (Figure S1A). Phylogenetically related non-parasitic species generally possess higher lncRNA counts compared to parasitic species (Figure S1A), suggesting that parasitism may influence lncRNA abundance.

Examining the lncRNA counts across different parasitic lifestyles, holoparasitic species exhibited the highest lncRNA counts, averaging 51% more than hemiparasites (Figure 1A; Figure S1B). The endoparasite *S. himalayana* showed a significant reduction, with only 1,507 lncRNAs identified (Figure 1A; Figure S1B). This reduction persisted even after normalization by genome size, establishing the trend: holoparasites > hemiparasites > endoparasites (Figure S1C). These findings suggest an association between parasitic lifestyle and lncRNA abundance, with holoparasites having notably higher counts, while endoparasites like *Sapria* exhibit a significant reduction despite their large genomes. To improve accuracy, high-confidence lncRNAs were also identified, providing a robust comparison across parasitic groups. While this refinement did not alter the observed trends (Figure 1A), focusing only on high-confidence lncRNAs reduced overall counts, potentially losing important information. Thus, subsequent analyses used the total number of identified lncRNAs to ensure a more comprehensive dataset.

**Figure 1.**
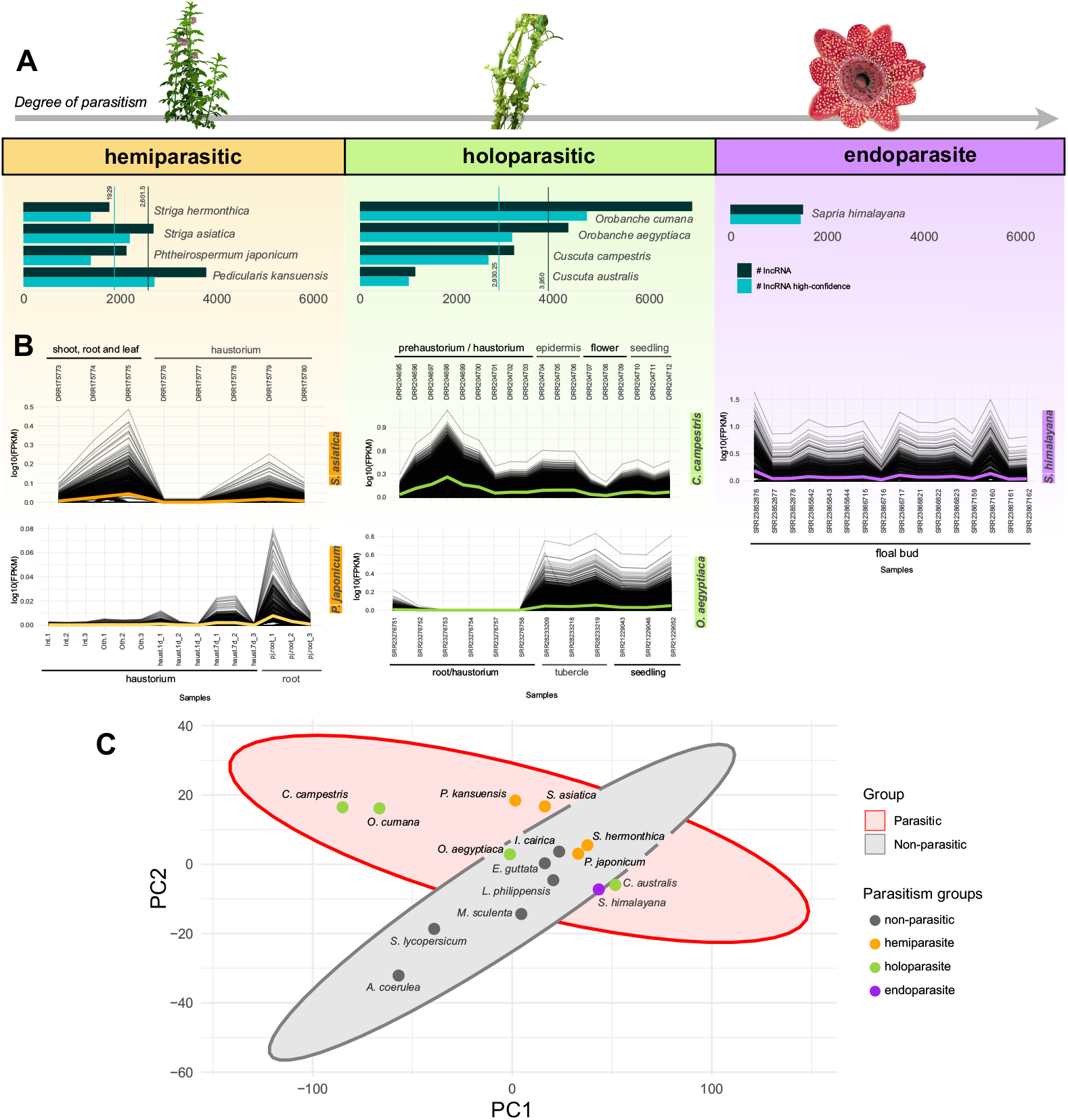
(A) Barplot showing the number of lncRNAs and high-confidence lncRNAs identified in this study across parasitic plant species with different levels of parasitism, (B) their expression profiles, and (C) PCA highlighting the distinct lncRNA characteristics compared to non-parasitic species.

To further explore the functional aspects of parasitic lifestyles, we analyzed lncRNA expression profiles using RNA-Seq data from public databases and examined qualitative characteristics such as k-mer composition, secondary structures, and sequence lengths across parasitic and non-parasitic groups. In hemiparasitic species like *Phtheirospermum japonicum*, lncRNA expression is predominantly observed in roots not actively interacting with a host (Figure 1B), suggesting its role is more associated with general plant development than parasitic interaction. Similarly, in *Striga asiatica*, higher expression levels were observed in the leaf, root, and shoot rather than in haustorial tissues (Figure 1A). In contrast, lncRNA expression in holoparasitic and endoparasitic species appears more balanced across organs but varies by species (Figure 1B). For example, in *Cuscuta*, expression is highest in haustorial tissues, indicating a strong link to haustorial function, while in *Orobanche*, expression is more prominent in non-haustorial tissues, suggesting additional roles. Interestingly, floral lncRNA expression is relatively low in both *Sapria* and *Cuscuta*, despite the reproductive importance of these tissues in their parasitic lifecycles. Furthermore, to investigate structural and sequence features, we performed a Principal Component Analysis (PCA), which revealed that lncRNAs in parasitic species exhibit fundamentally different characteristics compared to those in non-parasitic species, suggesting that parasitism directly influences lncRNA structural and compositional traits (Figure 1C).

To explore the possible origins of lncRNAs in parasitic plants, we analyzed repetitive motifs within these genomes, as it is well established that lncRNA origins are frequently associated with repetitive sequences across various groups of organisms^17^. We found no strong correlation between the overall percentage of repetitive elements and the total number of lncRNAs per genome in parasitic species (Figure S2A). Notably, *Sapria*, the species with the highest percentage of repetitive elements in its genome, exhibits a much lower lncRNA count compared to most other species (Figure S2A), indicating that lncRNA abundance is not simply a function of total repetitive element content. When examining specific types of repetitive elements, however, we observed a strong positive correlation between the number of lncRNAs and the percentage of LTR (long terminal repeat) elements in the genomes of parasitic plants (Figure S2B). This association suggests that LTR elements may play a significant role in the origination or regulation of lncRNAs within these species. Conversely, the correlation between lncRNA abundance and the percentage of non-LTR elements or DNA transposable elements was negative (Figure S2B).

Parasitic plant genomes, especially holo- and endoparasites, have acquired significant host-derived sequences through horizontal gene transfer (HGT). For example, Sapria has approximately 1.2% of its genome derived from HGT, and species such as *Cuscuta, Orobanche*, and *Pedicularis kansuensis* also harbor host-transferred genes^18–20^. To investigate whether lncRNAs in parasitic plants may originate from similar transfers, we performed phylogenetic analyses on the lncRNA sequences identified in this study. We conducted BLAST analyses to identify hits against a curated dataset (Table S 4) that included host species and phylogenetically related non-host species. The majority of positive hits were observed in holoparasitic species. In Figure 2A, a heatmap illustrates the distribution of hits by species, revealing a higher number of host-specific matches in species such as *Striga, Sapria*, and *Cuscuta*. For example, Orobanchaceae species showed significant hits against Fabaceae e Curcubitales species, Sapria against *Tetrastigma voinierianum*, and *Cuscuta* against various species of Rosids but in low numbers. (Figure 2A). Using these positive hits from the heatmap, we constructed a phylogeny that revealed a substantial number of clades with potential HGT events in *Sapria* (Figure 2B). Interestingly, *Cuscuta australis* and *Striga asiatica* also displayed possible HGT events (Figure S3A-B), despite the lack of prior reports in the literature suggesting horizontal transfers in this species. These findings highlight the potential role of HGT in shaping the lncRNA landscape of parasitic plants, particularly in endoparasitic species.

**Figure 2.**
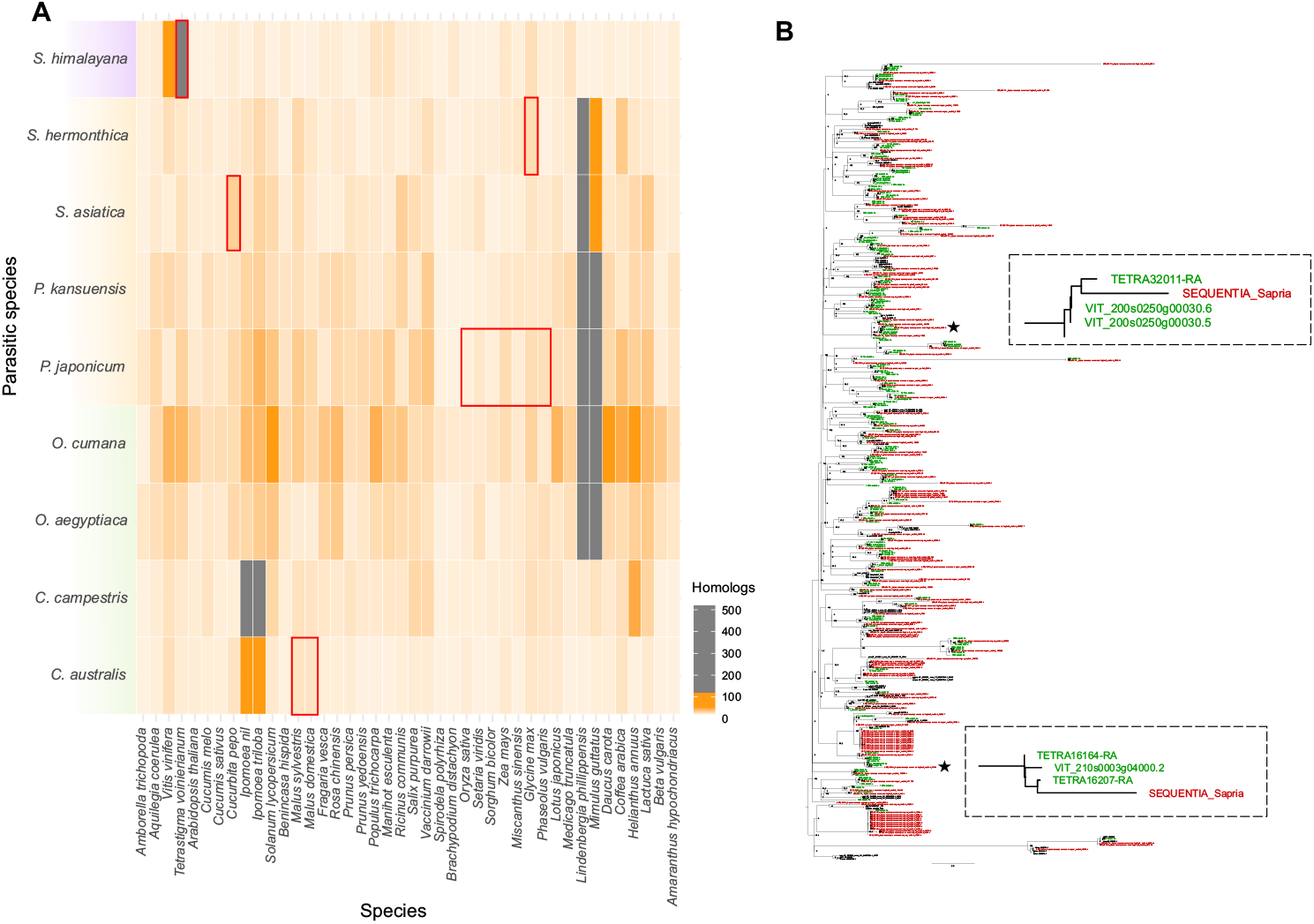
(A) Heatmap highlighting (red line box) homologous lncRNAs in distantly related parasitic plant species identified through BLAST and (B) phylogenetic tree indicating potential HGT events in *Sapria himalayana*.

Our findings provide new insights into the diversity, expression, and potential origins of lncRNAs in parasitic plants, highlighting their functional and evolutionary significance across different parasitic lifestyles. The distinct patterns of lncRNA abundance, structure, and expression between hemi-, holo-, and endoparasites suggest that these molecules are intricately linked to the unique biological and developmental adaptations of parasitism. The strong association between lncRNAs and key parasitic structures, such as haustoria, along with their presence in non-parasitic organs, underscores their multifunctional roles in both parasitism and general plant development. Furthermore, the evidence of horizontal transfer of lncRNA sequences, particularly in *Sapria*, opens new avenues for understanding how host-parasite interactions shape genomic innovation in parasitic plants. Together, these results highlight the importance of lncRNAs as central players in the molecular mechanisms underlying parasitism, providing a foundation for further exploration of their regulatory roles and evolutionary origins.

## Acknowledgements

This work was supported by the Serrapilheira Institute (grant number Serra-181226691) and by the Fundação de Amparo à Pesquisa do Estado de Minas Gerais (FAPEMIG) – Finance Code 001, that provided a PhD scholarship to WFCR.

## References

1. Kang, C. & Liu, Z. Global identification and analysis of long non-coding RNAs in diploid strawberry Fragaria vesca during flower and fruit development. BMC Genomics 16, 815 (2015).

2. Kitagawa, M., Kitagawa, K., Kotake, Y., Niida, H. & Ohhata, T. Cell cycle regulation by long non-coding RNAs. Cell. Mol. Life Sci. 70, 4785–4794 (2013).

3. Rai, M. I., Alam, M., Lightfoot, D. A., Gurha, P. & Afzal, A. J. Classification and experimental identification of plant long non-coding RNAs. Genomics 111, 997–1005 (2019).

4. Chen, Q. et al. From “Dark Matter” to “Star”: Insight Into the Regulation Mechanisms of Plant Functional Long Non-Coding RNAs. Front. Plant Sci. 12, 650926 (2021).

5. Song, L., Fang, Y., Chen, L., Wang, J. & Chen, X. Role of non-coding RNAs in plant immunity. Plant Communications 2, 100180 (2021).

6. Parasitic Plants. (Chapman & Hall, London Weinheim, 1995).

7. Press, M. C. & Phoenix, G. K. Impacts of parasitic plants on natural communities. New Phytologist 166, 737–751 (2005).

8. Khoei, M. A., Karimi, M., Karamian, R., Amini, S. & Soorni, A. Identification of the Complex Interplay Between Nematode-Related lncRNAs and Their Target Genes in Glycine max L. Front. Plant Sci. 12, 779597 (2021).

9. Han, G. et al. Identification of Long Non-Coding RNAs and the Regulatory Network Responsive to Arbuscular Mycorrhizal Fungi Colonization in Maize Roots. IJMS 20, 4491 (2019).

10. Yoshida, S., Cui, S., Ichihashi, Y. & Shirasu, K. The Haustorium, a Specialized Invasive Organ in Parasitic Plants. Annu. Rev. Plant Biol. 67, 643–667 (2016).

11. Yoshida, S. & Shirasu, K. Plants that attack plants: molecular elucidation of plant parasitism. Current Opinion in Plant Biology 15, 708–713 (2012).

12. Barkman, T. J. et al. Reading between the vines: Hosts as islands for extreme holoparasitic plants. American J of Botany 104, 1382–1389 (2017).

13. Shen, G. et al. Cuscuta australis (dodder) parasite eavesdrops on the host plants’ FT signals to flower. Proc. Natl. Acad. Sci. U.S.A. 117, 23125–23130 (2020).

14. Cai, L. et al. Deeply Altered Genome Architecture in the Endoparasitic Flowering Plant Sapria himalayana Griff. (Rafflesiaceae). Current Biology 31, 1002–1011.e9 (2021).

15. Sun, G. et al. Large-scale gene losses underlie the genome evolution of parasitic plant Cuscuta australis. Nat Commun 9, 2683 (2018).

16. Wu, Y. et al. Bidirectional lncRNA Transfer between Cuscuta Parasites and Their Host Plant. IJMS 23, 561 (2022).

17. Lee, H., Zhang, Z. & Krause, H. M. Long Noncoding RNAs and Repetitive Elements: Junk or Intimate Evolutionary Partners? Trends in Genetics 35, 892–902 (2019).

18. Lin, Q., Banerjee, A. & Stefanović, S. Mitochondrial Phylogenomics of Cuscuta (Convolvulaceae) Reveals a Potentially Functional Horizontal Gene Transfer from the Host. Genome Biology and Evolution 14, evac091 (2022).

19. Kado, T. & Innan, H. Horizontal Gene Transfer in Five Parasite Plant Species in Orobanchaceae. Genome Biology and Evolution 10, 3196–3210 (2018).

20. Fang, L. et al. Chromosome-level genome assembly of Pedicularis kansuensis illuminates genome evolution of facultative parasitic plant. Molecular Ecology Resources 24, e13966 (2024).

