## Supplemental Figure for "Investigating lncRNAs at different levels of parasitism in parasitic plants"

### Methods

#### lncRNA Identification

Putative lncRNAs were identified using predicted transcriptomes from nine parasitic plant species representing hemi-, holo-, and endoparasitic lifestyles (Table S1). First, transcripts longer than 200 nucleotides were filtered using a custom script, followed by filtering for transcripts with open reading frames (ORFs) shorter than 120 amino acids, a common characteristic of lncRNAs. After the initial filtering, BLASTx[1] searches were conducted against the SwissProt database (e-value < 0.001) to exclude protein-coding transcripts. Additionally, the Coding Potential Calculator (CPC2, https://cpc2.gao-lab.org/)[2] was employed to further select only non-coding transcripts, resulting in a preliminary list of potential lncRNAs for each species.

Next, this list was subjected to additional filtering steps. BLASTn (e-value < 0.05) was performed against a database of plant mature miRNA sequences from miRBase. The CMSCAN tool[3] was used to query the Rfam database, and MIReNA[4] was employed to identify and exclude precursor transcripts for small RNAs (sRNAs). This series of steps produced a final high-confidence set of lncRNAs for each species. Information regarding repetitive elements in the genomes of these species was extracted from the original genome sequencing publications (Tables S2).

#### Expression and PCA Analysis

Principal Component Analysis (PCA) was conducted using Jellyfish v2.3[5] to count k-mers and custom scripts to extract nucleotide lengths of the previously identified lncRNAs. Expression analyses were performed using RNA-Seq data available in the Sequence Read Archive (SRA) of NCBI (Table S3). After downloading the data, Salmon[6] v1.10.3 was used to quantify the expression levels of lncRNA transcripts. Visualization of the PCA and expression data was performed using R 4.4.0 and RStudio[7].

#### HGT Analysis

For the horizontal gene transfer (HGT) analysis, parasitic plant species with positive hits were identified using BLASTn (e-value < 1e-15) against a database constructed with transcriptomes of potential host plant species and phylogenetically related non-host species to enable clustering in phylogenetic analyses (Table S3). To ensure data accuracy and reduce redundancy, sequences with 100% identity were merged, and isoforms were removed using the SkipRedundant tool from EMBOSS v6.6.0.0 [8].

lncRNA transcripts were aligned using MAFFT[9] v7.490 with the following parameters: --thread 30, --reorder, --leavegappyregion, --maxiterate 1000, --retree 1, and --localpair. Phylogenetic trees were constructed using IQ-TREE [10] v2.0, applying ModelFinder[11] to identify the best substitution model for each family. Branch support was calculated using SH-aLRT with 1000 pseudoreplicates. The resulting phylogenetic trees were visualized using FigTree v1.4.4 (http://tree.bio.ed.ac.uk/software/figtree/).

### Supplemental Figures

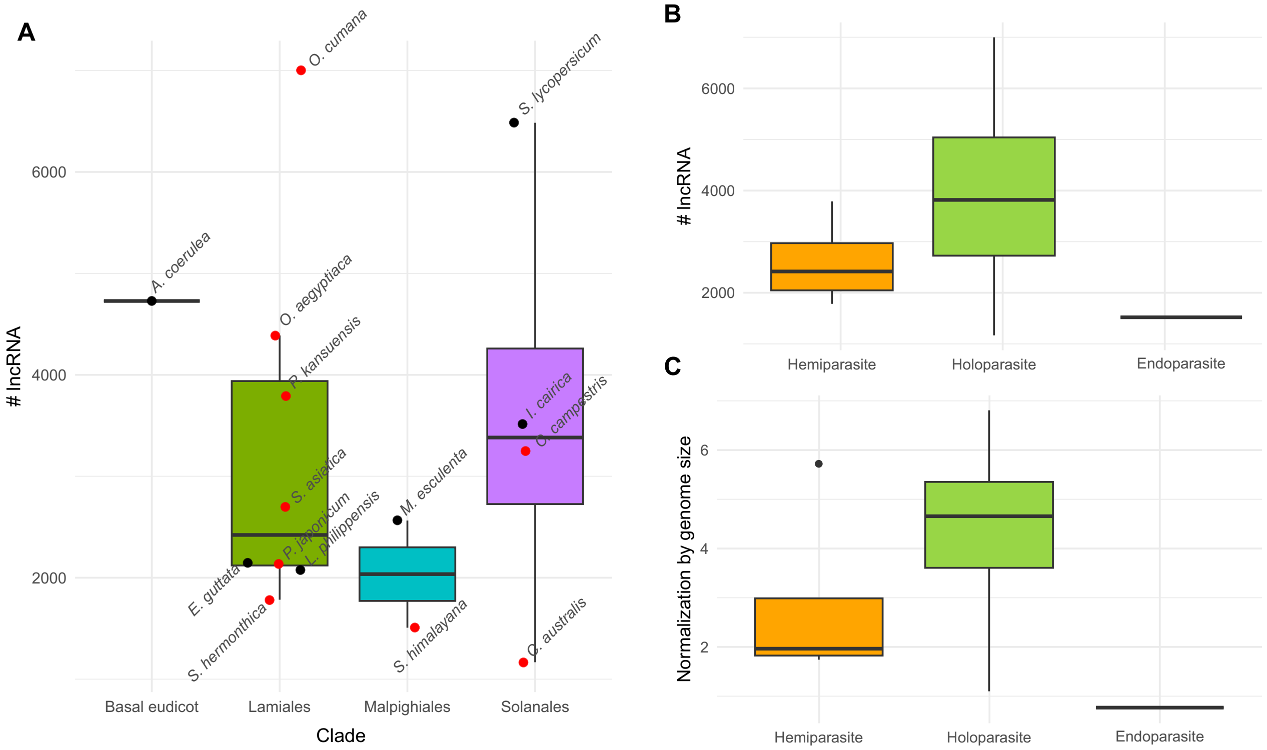


**Supplemental Figure 1.** Boxplots showing (A) the distribution of the number of lncRNAs identified in parasitic and phylogenetically related non-parasitic plants, (B) grouped by different levels of parasitism, and (C) normalized by genome size.

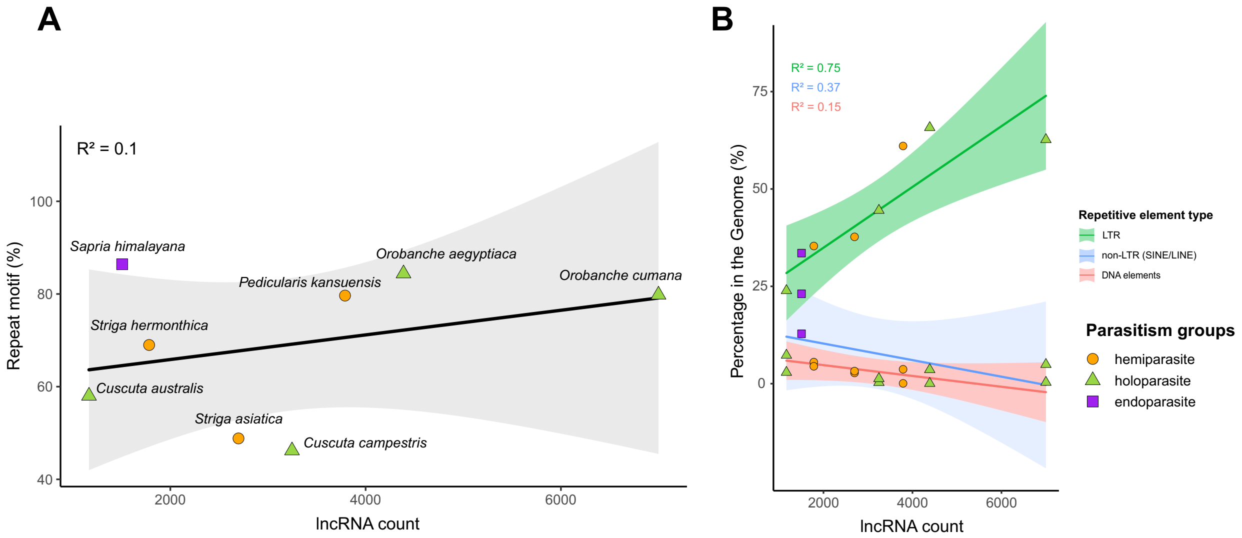


**Supplemental Figure 2.** Correlation plots between the number of lncRNAs and (A) the percentage of repetitive motifs and (B) different types of repetitive elements in the genomes of parasitic plant species.

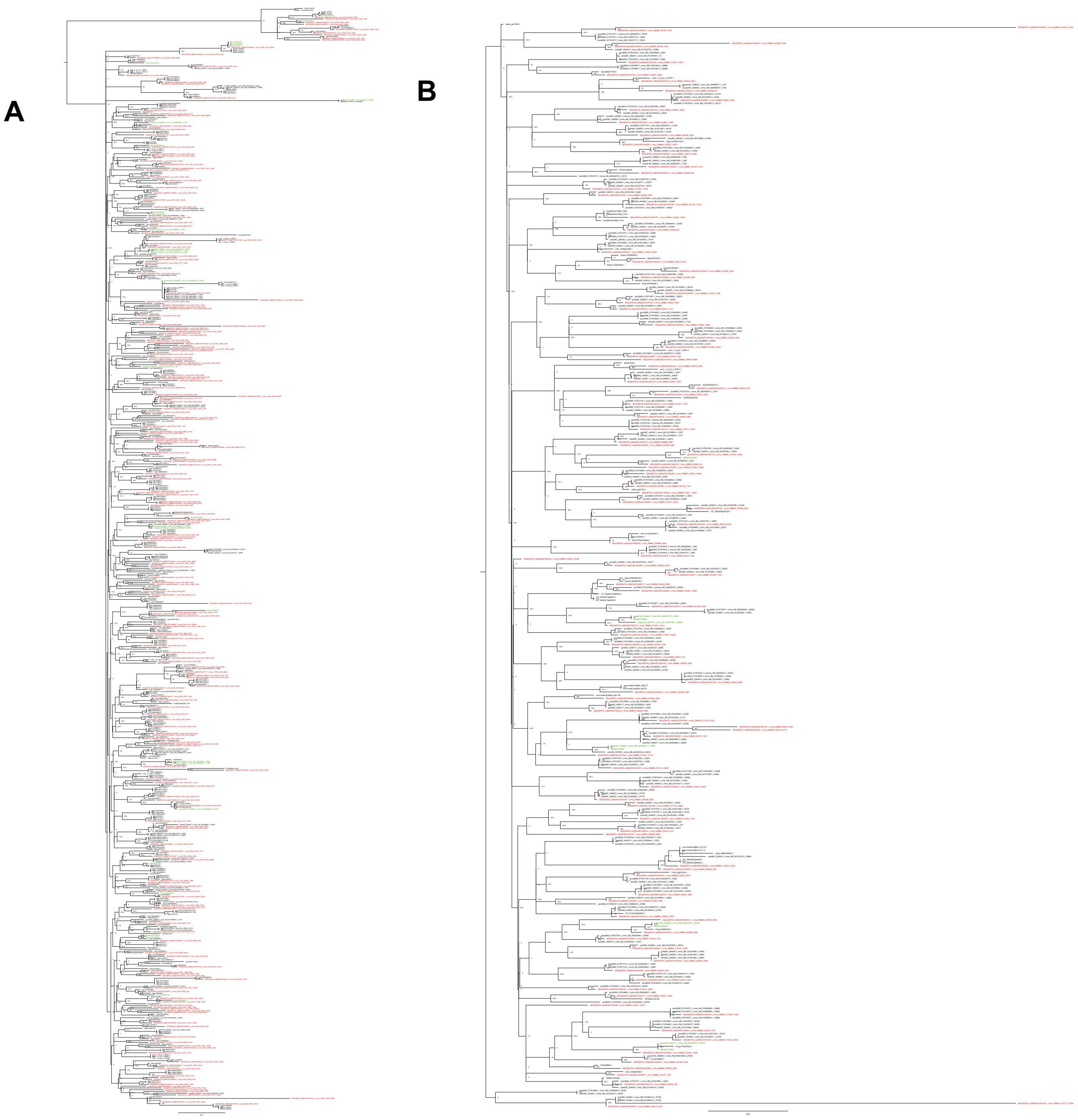


**Supplemental Figure 3.** Complete phylogenetic tree of (A) Striga asiatica and (B) Cuscuta australis, highlighting potential HGT events between the parasitic plants and distantly related species.

### References


1. Altschul, S. Gapped BLAST and PSI-BLAST: a new generation of protein database search programs. *Nucleic Acids Research* **25**, 3389–3402 (1997).
2. Kang, Y.-J. *et al.* CPC2: a fast and accurate coding potential calculator based on sequence intrinsic features. *Nucleic Acids Research* **45**, W12–W16 (2017).
3. Nawrocki, E. P. & Eddy, S. R. Infernal 1.1: 100-fold faster RNA homology searches. *Bioinformatics* **29**, 2933–2935 (2013).
4. Mathelier, A. & Carbone, A. MIReNA: finding microRNAs with high accuracy and no learning at genome scale and from deep sequencing data. *Bioinformatics* **26**, 2226–2234 (2010).
5. Marçais, G. & Kingsford, C. A fast, lock-free approach for efficient parallel counting of occurrences of *k* -mers. *Bioinformatics* **27**, 764–770 (2011).
6. Patro, R., Duggal, G., Love, M. I., Irizarry, R. A. & Kingsford, C. Salmon provides fast and bias-aware quantification of transcript expression. *Nat Methods* **14**, 417–419 (2017).
7. RStudio Team. *RStudio: Integrated Development Environment for R*. (RStudio, PBC., Boston, MA, 2020).
8. Rice, P., Longden, I. & Bleasby, A. EMBOSS: The European Molecular Biology Open Software Suite. *Trends in Genetics* **16**, 276–277 (2000).
9. Katoh, K. MAFFT: a novel method for rapid multiple sequence alignment based on fast Fourier transform. *Nucleic Acids Research* **30**, 3059–3066 (2002).
10. Nguyen, L.-T., Schmidt, H. A., von Haeseler, A. & Minh, B. Q. IQ-TREE: A Fast and Effective Stochastic Algorithm for Estimating Maximum-Likelihood Phylogenies. *Molecular Biology and Evolution* **32**, 268–274 (2015).
11. Kalyaanamoorthy, S., Minh, B. Q., Wong, T. K. F., von Haeseler, A. & Jermiin, L. S. ModelFinder: fast model selection for accurate phylogenetic estimates. *Nat Methods* **14**, 587–589 (2017).
